# The branched chain aminotransferase IlvE promotes growth, stress resistance and pathogenesis of *Listeria monocytogenes*

**DOI:** 10.1101/2020.01.31.929828

**Authors:** Karla D. Passalacqua, Tianhui Zhou, Tracy A. Washington, Basel H. Abuaita, Abraham L. Sonenshein, Mary X.D. O’Riordan

**Affiliations:** University of Michigan Medical School, Department of Microbiology & Immunology, Ann Arbor, MI; Tufts University School of Medicine, Department of Molecular Biology & Microbiology, Boston, MA

**Author notes:** Karla D. Passalacqua and Tianhui Zhou contributed equally to this work. Author order was determined on the basis of seniority.

## Abstract

The bacterial plasma membrane is a key interface during pathogen-host interactions, and membrane composition enhances resistance against host antimicrobial defenses. Branched chain fatty acids (BCFAs) are the major plasma membrane component in the intracellular Gram-positive pathogen *Listeria monocytogenes* (Lm) and BCFA metabolism is essential for Lm growth and virulence. BCFA synthesis requires branched chain amino acids (BCAAs), and the BCAA Isoleucine (Ile) is a necessary substrate for the predominant membrane anteiso-BCFAs (ai-BCFAs) as well as an environmental signal for virulence regulation in Lm. In this study, we explored how two proteins that metabolize or sense Ile contribute to Lm growth, BCFA metabolism, and virulence. The IlvE aminotransferase incorporates Ile into ai-BCFAs, while CodY is an Ile-sensing regulator that coordinates BCAA synthesis and virulence gene expression. Analysis of deletion mutants lacking IlvE (Δ*ilvE*) or CodY (Δ*codY*) revealed a major role for IlvE under nutrient restriction and stress conditions. Cultures of the Δ*ilvE* mutant contained proportionally less ai-BCFAs relative to wild type, while of the Δ*codY* mutant had a lower proportion of ai-BCFAs in stationary phase, despite containing more cell-associated Ile. Both Δ*ilvE* and Δ*codY* mutants required exogenous Ile for optimal growth, but the Δ*ilvE* mutant had an absolute requirement for Valine and Leucine when Ile was absent. IlvE was also necessary for resistance to membrane stress, cell-to-cell spread, infection of primary macrophages, and virulence in mice. Our findings implicate IlvE as an integral aspect of Lm stress resistance and emphasize the central importance of Ile in Lm growth and virulence.

## INTRODUCTION

The bacterial plasma membrane is a key interface of pathogen-host interactions and an important intrinsic barrier to host antimicrobial defenses. Situated just beneath and intimately connected to the bacterial cell wall, the plasma membrane is a crucial structure of the bacterial cell surface; thus, the interface between bacterium and host cell is of particular importance for intracellular pathogens such as *Listeria monocytogenes* (Lm), the causative agent of listeriosis (1–3). During its infectious cycle, Lm enters mammalian cells and traverses through a range of cellular locales, each with distinct nutrient availability, redox state, and host antibacterial mechanisms (4). Here, the bacterial membrane serves as an environmental sensor and a defensive structure central to intracellular survival and replication (5). Therefore, exploring Lm membrane dynamics is central for elucidating virulence strategies of this important pathogen.

As in many Gram-positive bacteria, including *Staphylococcus aureus*, the Lm plasma membrane is predominantly composed of branched chain fatty acids (BCFAs), a structural feature important for bacterial integrity against multiple stresses and during pathogenesis (6–14). Odd numbered (C15, C17) anteiso-BCFAs (ai-BCFAs) are the most abundant form of BCFA in the Lm membrane, and the ability of Lm to thrive in cold temperatures is due in large part to high ai-BCFA content that enhances membrane fluidity (15–17). To optimize membrane fluidity in different environments, Gram-positive bacteria alter the ratio of ai-BCFAs to iso-BCFAs, where ai-BCFAs contribute to higher fluidity due to positioning of the terminal methyl groups on the acyl chains (18). Because BCFA synthesis depends on the acquisition and/or biosynthesis of branched chain amino acids (BCAAs: Isoleucine, Leucine and Valine [Ile, Leu, Val]) (7, 19), membrane remodeling and BCAA metabolism are tightly linked. While Lm is fully capable of synthesizing BCAAs *de novo*, exogenous BCAAs are required for optimal growth due in part to high demand for BCFAs in the membrane and the activity of a ribosome-mediated attenuator that limits BCAA synthesis (20, 21). During infection, Lm replicates inside host cells where BCAAs and other nutrients are limited and may be actively withheld from bacteria by host defense mechanisms (22, 23). Therefore, the ability of Lm to acquire host BCAAs and to make *de novo* BCAAs, especially Ile, to generate membranes with high BCFA levels is critical to Lm pathogenesis.

Branched chain amino acid aminotransferase (BCAT) enzymes initiate bacterial BCFA synthesis by converting BCAAs into branched chain α-keto acids. Downstream of BCAT, branched chain α-keto dehydrogenase enzymes (BKD) produce acyl coenzyme A (CoA) molecules that are the primers for fatty acid synthesis (Fig. 1A) (9). While BKD is essential for BCFA metabolism and protection from host immune defenses such as antimicrobial peptides (12, 13), the Lm BCAT IlvE is required for resistance to the compound *trans*-cinnamaldehyde, a small molecule with anti-microbial properties (14, 24). Additionally, the transcriptional regulator CodY, which senses BCAA and GTP levels, plays a major role in coordinating BCAA metabolism with virulence gene expression (21, 25–28). Importantly, when Ile levels are high, CodY inhibits *de novo* BCAA synthesis, and when Ile levels are low, this inhibition is relieved, allowing the bacteria to synthesize vital BCAAs (7). Thus, the ability of Lm to sense and regulate BCAA levels, particularly Ile, and to implement BCFA remodeling is an important attribute for adaptation to changing environments, especially in stress conditions as found in the mammalian host.

**Figure 1.**
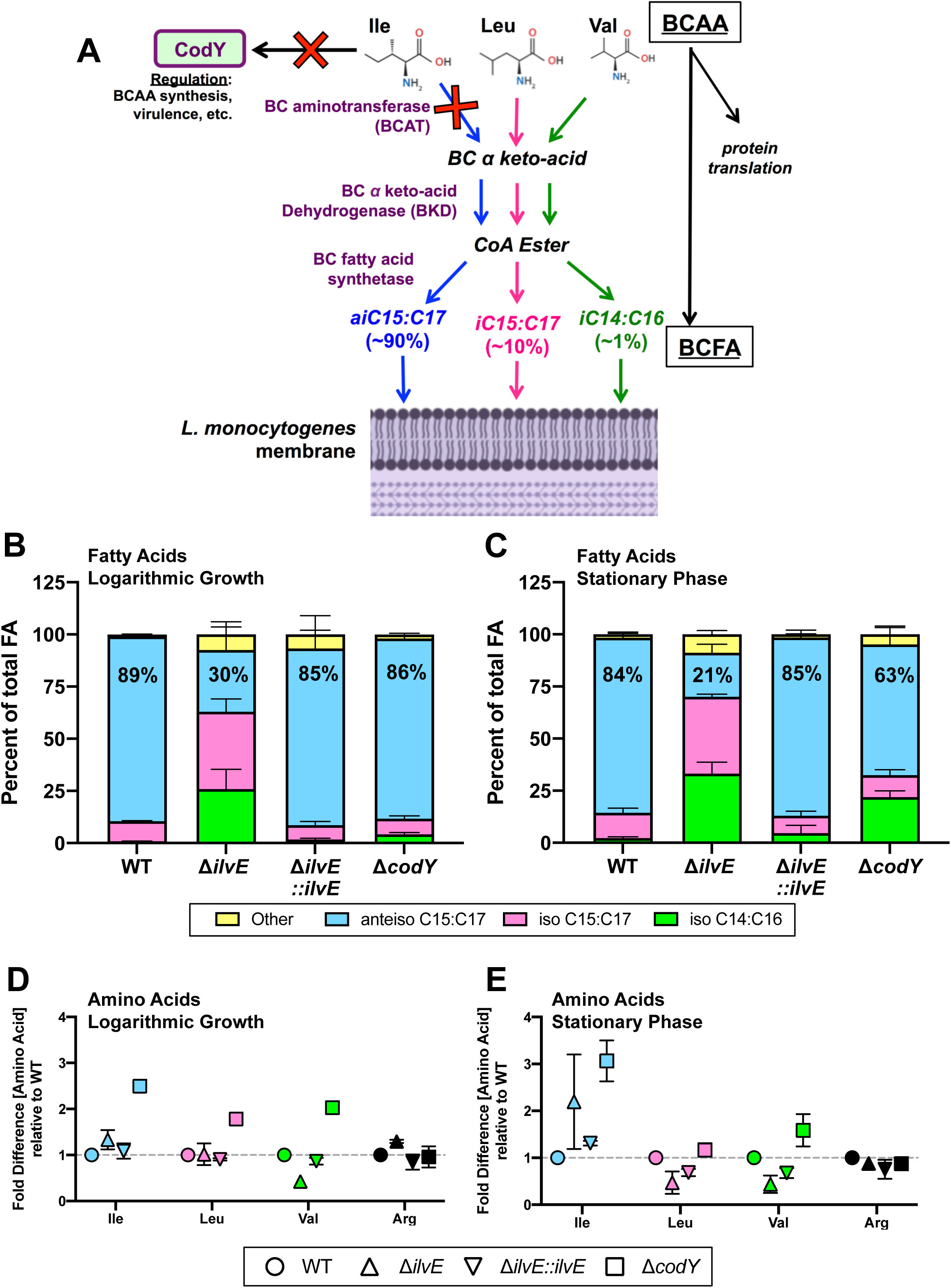
Changes in Fatty Acid and BCAA content in Lm lacking IlvE or CodY. (A) Simplified overview of branched chain fatty acid (BCFA) biosynthesis in Gram-positive bacteria (based on detailed diagram in (9)) showing pathways that incorporate branched chain amino acids (BCAAs: Ile, Leu & Val). Red X represents points in pathways where deletion mutants were used in this study. Colored arrows indicate pathways of individual BCAAs that are incorporated into final BCFA isoforms (18). Purple text = enzyme names. (B and C) Graphs represent the relative amounts of the major fatty acids as a percentage of total fatty acids contained in Lm cultures of WT, Δ*ilvE*, Δ*ilvE::ilvE^+^* and Δ*codY* strains grown in nutrient limiting medium (LDM) to (B) mid-logarithmic and (C) stationary phase. Graphs represent combined data from three independent experiments. Graphs shown here and data in Tables 2 and 3 are the combined quantities of odd numbered (C15 and C17) or even-numbered (C14 and C16) BCFAs. Individual numbered species (e.g., ai-C15 only) and all other fatty acids are in Supplemental Tables S3 and S4. (D and E). Cultures of WT, Δ*ilvE*, Δ*ilvE::ilvE* and Δ*codY* strains grown in LDM to (D) mid-logarithmic and (E) stationary phase were analyzed by mass spectrometry. Concentrations of BCAAs were normalized to total protein content and are shown as ratios relative to WT. Error bars show the range of fold difference compiled from 2 independent experiments.

Due to the central requirement for Ile in promoting membrane integrity through generation of ai-BCFAs and for engaging CodY regulatory activity, we hypothesized that proteins involved in Ile metabolism are central to the ability of Lm to cause disease. Therefore, we used a genetic approach to explore how the BCAT IlvE and the regulator CodY contribute to membrane dynamics, growth and pathogenesis of Lm. Here we show that deficiency of either IlvE or CodY can alter membrane fatty acid content, but bacteria lacking IlvE are very susceptible to membrane stress and nutrient limitation and are less fit in *in vitro* and *in vivo* infection models.

## RESULTS

### Membrane anteiso-BCFA generation requires IlvE and relies on CodY for homeostasis in stationary phase in nutrient restricted medium

Isoleucine (Ile) is an essential metabolite for protein translation and for synthesis of the high ai-BCFA membrane content of Lm (Fig. 1A), and the aminotransferase IlvE is predicted to be the first enzyme that commits Ile into the biosynthetic pathway for odd-numbered (C15, C17) ai-BCFAs. Low availability of Ile in the intracellular environment during infection is thought to act as a signal for Lm to coordinate metabolism and virulence, mainly through the Ile-sensing transcriptional regulator CodY (21, 26, 27). To characterize dynamics of Ile usage in BCFA biosynthesis and virulence, we assessed deletion mutant strains lacking IlvE or CodY (Δ*ilvE* and Δ*codY* mutants) (Table S1 and Methods).

**Table 1.** Fatty Acid Content of L. monocytogenes during Logarithmic Growth in LDM

|  | Percent of Total Fatty Acid Content – Logarithmic Growth |  |  |  |
| --- | --- | --- | --- | --- |
|  | Mean Percent (SD) |  |  |  |
| | WT | $\Delta ilvE$ | $\Delta ilvE::ilvE^+$ | $\Delta codY$ |
| **Other** | 0.95 (0.20) | 7.41 (3.58) | 6.72 (9.05) | 1.91 (0.54) |
| **anteisoC15:C17** | 88.56 (0.30) | ****29.67 (13.54) | 84.67 (8.76) | 86.41 (1.44) |
| **isoC15:C17** | 9.60 (0.30) | ****37.06 (6.15) | 6.85 (1.80) | 7.43 (1.36) |
| **isoC14:C16** | 0.89 (0.11) | ****25.86 (9.50) | 1.76 (0.61) | 4.24 (0.86) |

Two-Way ANOVA using Dunnett's Multiple Comparisons Test (compare rows within columns) compared to Wild Type (WT). \*\*\*\* $P < 0.0001$ .

Previously, an *ilvE* transposon-generated null mutant was shown to have extremely low levels of ai-BCFAs when grown in rich, undefined BHI medium (14). Therefore, we predicted that the Δ*ilvE* strain created for this study would have substantially lower levels of ai-BCFA when grown in a nutrient-limited medium (Fig. 1A). Additionally, we predicted that the Δ*codY* mutant would generate ai-BCFA levels equivalent to WT, since eliminating CodY inhibition of *de novo* BCAA synthesis should increase bacterial BCAA levels, resulting in ample building blocks for ai-BCFAs. Note that recent RNAseq analysis showed that the expression of the Ile, Leu, Val-production operon increased substantially in the Δ*codY* mutant in both rich medium (BHI) and in nutrient-limited medium (29), but not when BCAAs were extremely limited; moreover, the *codY* null mutation had no significant impact on *ilvE* transcription in any media tested (T. A. Washington, A. L. Sonenshein and B. R. Belitsky, manuscript in preparation) despite the fact that CodY has a relatively strong binding site upstream of *ilvE* (30), which could also be a regulatory binding site for the locus upstream of *ilvE*. Therefore, to test the role of IlvE and CodY in fatty acid metabolism, we measured total fatty acid content in WT, Δ*ilvE*, Δ*ilvE::ilvE^+^* (*ilvE* complemented) and Δ*codY* strains grown to mid-logarithmic and stationary phase in *Listeria* defined medium (LDM - contains seven amino acids including BCAAs) (29), a nutrient-limited medium (Fig. 1B-C, Tables 1-2, Tables S3-S4).

**Table 2.** Fatty Acid Content of L. monocytogenes during Stationary Phase in LDM

|  | Percent of Total Fatty Acid Content – Stationary Phase |  |  |  |
| --- | --- | --- | --- | --- |
|  | Mean Percent (SD) |  |  |  |
| | WT | $\Delta ilvE$ | $\Delta ilvE::ilvE$ | $\Delta codY$ |
| **Other** | 1.65 (0.44) | *8.86 (1.85) | 1.53 (0.21) | 4.84 (3.27) |
| **anteisoC15:C17** | 83.87 (2.76) | ****21.06 (4.14) | 85.55 (3.53) | ****62.75 (8.89) |
| **isoC15:C17** | 12.26 (2.09) | ****36.87 (1.23) | 8.32 (2.23) | 10.44 (2.69) |
| **isoC14:C16** | 2.22 (0.66) | ****33.22 (5.51) | 4.60 (3.75) | ****21.97 (3.03) |

Two-Way ANOVA using Dunnett's Multiple Comparisons Test (compare rows within columns)
compared to Wild Type (WT). \* $P < 0.05$ ; \*\*\*\* $P < 0.0001$
Data are combined from n = 3 experiments

**Table 3.** ^1^Doubling times of *L. monocytogenes* strains in small volume growth analysis in rich (BHI) and nutrient-limiting (LDM) medium.

| | WT | $\Delta ilvE$ | $\Delta ilvE::ilvE^+$ | $\Delta codY$ |
| --- | --- | --- | --- | --- |
|  | Mean (SD) in minutes |  |  |  |
| **BHI** | 65.9 (5.9) | 71.0 (2.3) | 59.5 (18.1) | 63.8 (3.3) |
| **LDM** | 118.5 (5.9) | 196.1 (17.0) | 204.9 (13.9) | 110.9 (6.5) |

<sup>1</sup>Calculated per (40). Data are combined from n = 4 independent experiments combined

The Δ*ilvE* strain contained significantly lower proportions of ai-BCFAs in the total fatty pool compared to WT (Fig. 1B-C). During both culture phases, WT cells had greater than 80% ai-BCFAs, while Δ*ilvE* cells had 30% ai-BCFAs in logarithmic phase and 21% ai-BCFAs in stationary phase. Whereas WT cells had extremely low levels of iso-BCFAs (0.9 – 12%), the Δ*ilvE* strain included substantial levels odd-numbered iso-C15 and iso-C17 fatty acids (37%) and even-numbered iso-C14 and iso-C16 (26% to 33%) (Tables 1 and 2). This indicates that in the absence of IlvE, Lm must incorporate the other BCAAs (Leu and Val) into BCFAs. The complemented strain Δ*ilvE::ilvE^+^* showed an almost identical fatty acid profile to WT in both phases. Interestingly, the Δ*codY* mutant differed markedly from WT in stationary phase. Here, ai-BCFAs made up 63% of the fatty acid profile, and even-numbered iso-BCFAs increased to 22% (ten-fold higher than WT). These results suggest that CodY may contribute to membrane BCFA remodeling when salvageable nutrients are depleted, and may repress genes involved in iso-BCFA synthesis during stationary phase.

We conclude that IlvE is a major driver for ai-BCFA generation during Lm growth in nutrient-limited medium; however, bacteria lacking IlvE were still able to generate more than 20% ai-BCFAs, suggesting the presence of another aminotransferase that is able to incorporate Ile into ai-BCFA. Additionally, we conclude that CodY is involved in BCFA homeostasis during stationary phase in LDM.

### Bacteria lacking CodY harbor higher levels of BCAA compared to WT

CodY is a sensor of BCAAs in Gram-positive bacteria, particularly Ile, and controls BCAA synthesis when these important metabolites are at low levels (31, 32). We initially hypothesized that bacteria lacking CodY would constitutively synthesize BCAAs in addition to acquiring exogenous BCAAs, and therefore should be well positioned to generate sufficient levels of ai-BCFAs regardless of growth phase. However, the observation that the Δ*codY* mutant had lower levels of ai-BCFAs in stationary phase in LDM (Fig. 1C) prompted us to directly measure levels of cell-associated BCAAs during growth in LDM. We therefore grew WT, Δ*codY*, Δ*ilvE* and Δ*ilvE::ilvE^+^* strains in LDM to mid-logarithmic and stationary phase, removed the extracellular medium, and assessed cell-associated BCAA content by mass spectrometry (Fig. 1D-E). Unsurprisingly, Δ*codY* lysates contained higher levels of all three BCAAs relative to WT during logarithmic growth, with Ile being the highest (∼2.5-fold higher relative to WT), confirming the role of CodY for BCAA synthesis during nutrient-restriction. Notably, we observed approximately three-fold more Ile in stationary phase cultures for the Δ*codY* mutant relative to WT, but similar levels of Leu and Val. Thus, although the Δ*codY* strain in stationary phase contains more Ile available for ai-BCFA compared to WT, this strain does not match WT levels of Ile incorporation into ai-BCFAs. These data suggest that CodY may play a role in membrane ai-BCFA homeostasis during stationary phase through an as yet undefined mechanism.

Since bacteria lacking IlvE showed a severe reduction in ai-BCFA content in LDM, we hypothesized that the Δ*ilvE* mutant would harbor higher levels of cell-associated Ile during all growth phases in LDM compared to WT due to the lack of incorporation of this amino acid into ai-BCFAs. However, we observed similar levels of Ile in the Δ*ilvE* mutant and in WT during logarithmic growth, and wide variability of cell-associated Ile in the Δ*ilvE* strain during stationary phase (Fig. 1D-E). Also, Val was approximately half the level in the Δ*ilvE* mutant compared to WT in logarithmic and stationary phase (Fig. 1D-E), while the Leu level was less than half of WT only in stationary phase. These data suggest that when IlvE is lacking, Lm uses Val and Leu for BCFA metabolism.

### Growth in BCAA-limiting conditions requires branched-chain aminotransferase IlvE

While fatty acid analysis represents relative levels of lipid species in a population of cells, these data do not reveal differences in growth rate between strains. Therefore, we examined the contributions of IlvE and CodY to bacterial growth in nutrient replete Brain-Heart Infusion medium (BHI) and in nutrient-limited LDM. All growth experiments were initiated using bacteria grown to mid-logarithmic phase in LDM. We hypothesized that because the absence of CodY normally contributes to increased BCAA biosynthesis during Ile limitation (26), Δ*codY* bacteria would grow as well as, or better than, WT bacteria in LDM. We also hypothesized that growth of the Δ*ilvE* mutant would be slower than WT in nutrient-limited medium due to its severe reduction in ai-BCFAs, a major membrane component for Lm.

In BHI, both the Δ*ilvE* and the Δ*codY* mutants grew equivalently to WT, showing that IlvE and CodY are not essential for Lm growth in a nutrient rich environment (Fig. 2A and Table 3). In LDM containing all three BCAAs at 100 μg/mL, the Δ*ilvE* strain grew slightly more slowly than WT, whereas the Δ*codY* strain grew the same as, or slightly better than, WT (Fig. 2B). Although the Δ*ilvE* culture reached the same maximum density as WT in LDM, its doubling time during logarithmic growth was about 1.7-fold longer than WT (Table 3). These data reveal that ai-BCFA synthesis through IlvE contributes to bacterial growth rate when nutrients are limited. In LDM, the Δ*ilvE::ilvE^+^* complemented strain also grew more slowly than WT, despite the fact that it was able to generate BCFA profiles similar to WT in this medium (Fig. 1B-C). We therefore asked whether the *ilvE* gene is expressed at WT levels in the complemented strain. Indeed, RT-qPCR of *ilvE* expression revealed lower transcript levels of this gene in the complemented strain (Fig. 2C), particularly during exponential growth.

**Figure 2.**
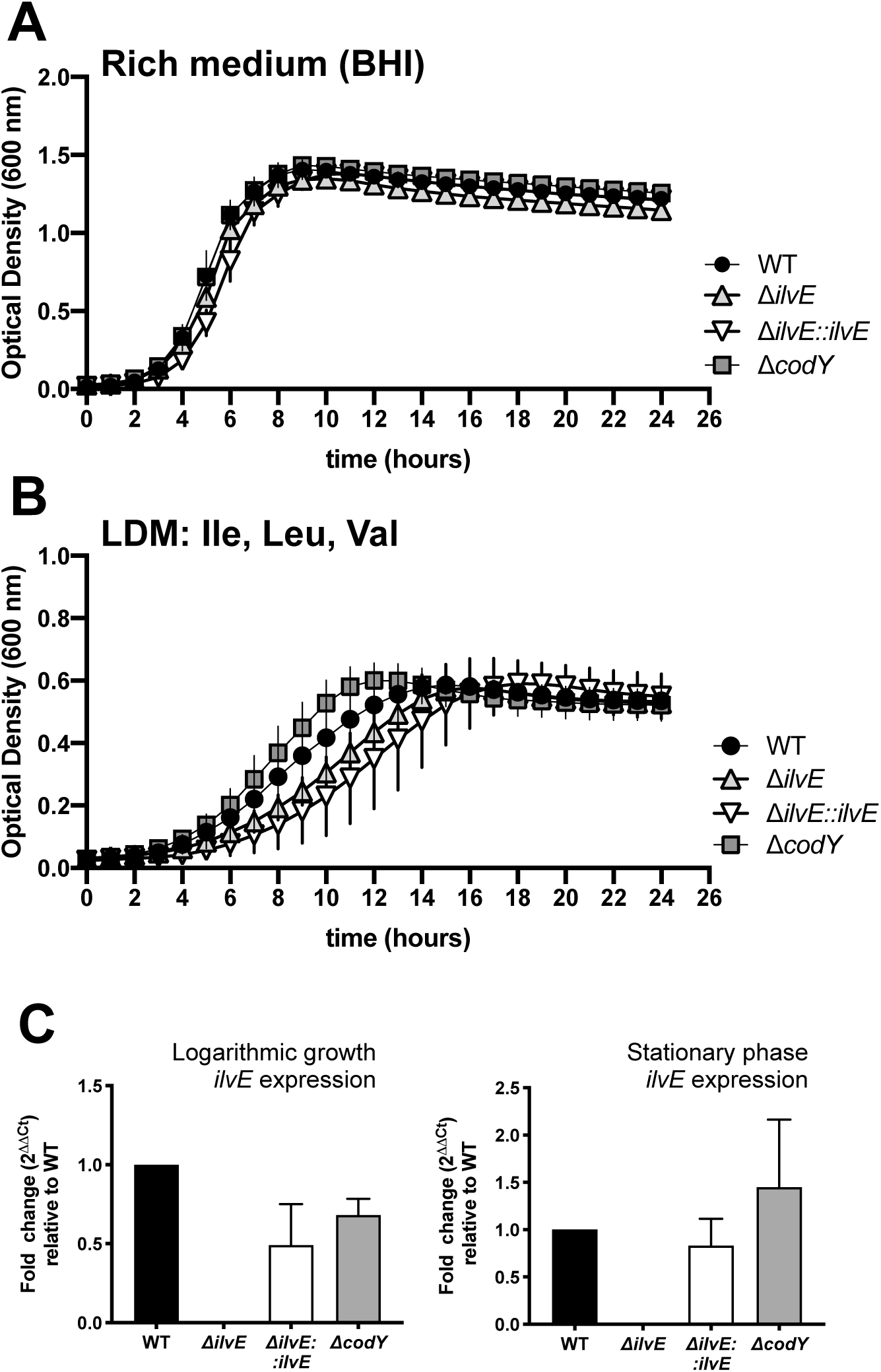
Growth of Δ*ilvE* and Δ*codY* mutants in rich and nutrient-limited medium. Bacterial growth of WT (circles), Δ*ilvE* (triangles), Δ*ilvE::ilvE^+^* (inverted triangles), and Δ*codY* (squares), was analyzed on a Bioscreen instrument. Samples were inoculated from recovered frozen cultures that had been prepared in LDM to mid-logarithmic phase. Optical Density at 600 nm (OD600) was measured for 24 hours at 37°C with shaking. Experiments include growth in (A) Rich medium = Brain Heart Infusion (BHI) and (B) LDM containing amino acids at 100 μg/mL. Data are compiled from three independent experiments with three technical replicates per experiment. Each point is the mean with error bars representing the Standard Deviation. (C) RT-qPCR analysis of *ilvE* expression in Lm grown in LDM to logarithmic (left) and stationary (right) phase.

Fatty acid distributions (Fig. 1B-C) suggested that mutants lacking IlvE or CodY use Val and Leu for synthesis of iso-BCFAs at higher levels than WT. Therefore, we asked how Δ*ilvE* and Δ*codY* strains would grow when one or all three of the BCAAs are lacking in the growth medium, despite having the ability to synthesize all three BCAAs *de novo*. In LDM lacking all three BCAAs, all four strains grew poorly, with WT and Δ*ilvE::ilvE^+^* reaching the highest optical density at 600 nm (OD600) of ∼0.4, compared to all strains reading ∼0.6 in LDM when all BCAAs were present (Fig. 2B versus Fig. 3A). Additionally, the Δ*ilvE* mutant exhibited large variability when no BCAAs were supplied, while Δ*codY* grew the poorest (Fig. 2B versus Fig. 3A). The fact that the Δ*codY* mutant grew so poorly in medium with no exogenous BCAAs was surprising considering that this mutant has no restriction on *de novo* synthesis of BCAAs. These data support the semi-auxotrophic nature of Lm for BCAAs, highlighting the importance of exogenous BCAAs for optimal growth and revealing a key role for IlvE and CodY when all exogenous BCAAs are unavailable.

**Figure 3.**
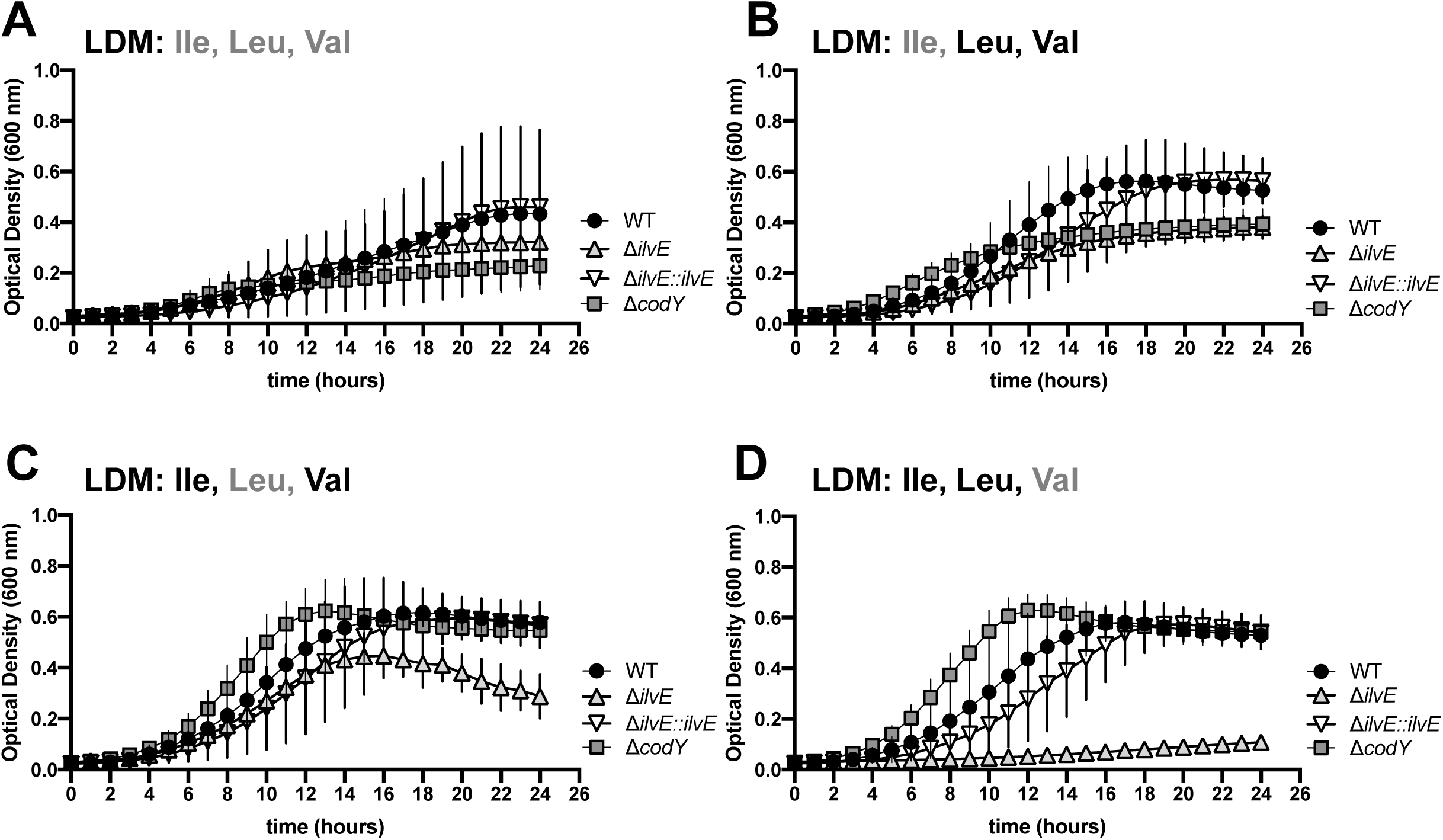
Growth of Lm in LDM with variable exogenous BCAAs. Bacterial growth of WT (circles), Δ*ilvE* (triangles), Δ*ilvE::ilvE^+^* (inverted triangles), and Δ*codY* (squares), performed as in Figure 2, but in LDM containing (A) no BCAAs, (B) no Ile (Val & Leu only), (C) no Leu (Ile & Val only), and (D) no Val (Ile & Leu only). Data are compiled from three independent experiments with three technical replicates per experiment. Each point is the mean with error bars representing standard deviation.

While IlvE is needed for enzymatic incorporation of Ile into BCFAs, CodY specifically senses and binds cellular Ile (33, 34). Due to their specific relationships with Ile, we then asked how Δ*ilvE* and Δ*codY* mutants would grow when exogenous Ile is lacking but when Val and Leu are present. Interestingly, both Δ*ilvE* and Δ*codY* strains were similarly attenuated when only Ile was lacking, reaching a lower maximum density compared to WT and Δ*ilvE::ilvE^+^* strains (Fig. 3B). Again, this was unexpected for the Δ*codY* mutant, since we predicted that the Δ*codY* strain would have no growth defect in the absence of Ile due to its higher cell associated Ile concentrations (Fig. 1D-E). These results reveal a complex role for CodY in Ile sensing and BCAA homeostasis. We conclude that both IlvE and CodY are required for optimal bacterial growth when exogenous Ile is absent.

When either all three BCAAs or only Ile were absent in the growth medium (Fig. 3A-B), the Δ*ilvE* and Δ*codY* mutants were attenuated for growth to a similar degree. However, in media containing exogenous Ile but lacking either of the other two BCAAs (Leu or Val), the two mutants revealed unique growth phenotypes (Fig. 3C-D). In LDM containing Ile and one other BCAA (Val or Leu), the Δ*codY* mutant grew more robustly than WT, suggesting a dominant role for Ile in Lm growth when CodY regulation is lacking. But the Δ*ilvE* mutant showed strict requirements for Leu and Val in the presence of Ile. When only Leu was absent (ie, Ile and Val present), the Δ*ilvE* mutant grew as it did in normal LDM (with all BCAAs, Fig. 2B) for about 12 hours, but reached stationary phase early and then had a decrease in OD600 (Fig. 3C). When only Val was absent (ie, Ile and Leu present), the Δ*ilvE* strain was entirely unable to grow (Fig. 3D), revealing an absolute requirement for Val when exogenous Ile is not incorporated into ai-BCFAs by IlvE. The IlvE complemented strain was able to eventually reach a maximum density in stationary phase similar to that of WT in all of these conditions (Fig. 3A-D), albeit at a slightly slower rate. Thus, when IlvE is not present, Lm has an increased dependence on Val and Leu for growth. These data show that while CodY is tightly linked to Ile sensing and homeostasis, IlvE activity plays a key role in cellular homeostasis when any of the individual BCAAs are lacking exogenously. Collectively, these growth trends indicate that BCAA levels are controlled at multiple levels in Lm.

### *Listeria* lacking IlvE exhibit decreased intracellular replication in macrophages and reduced cell-to-cell spread

Having established that IlvE and CodY play a role in generating membrane ai-BCFAs and in promoting optimal growth in BCAA-limited environments, we asked whether these proteins specifically contribute to Lm pathogenesis. We hypothesized that the Δ*ilvE* mutant would be less efficient at intracellular growth in a cell culture infection model due to its relatively slow growth during nutrient restriction. Previously, the Δ*codY* mutant strain has shown different behaviors in various *in vitro* macrophage infections models (25, 28). Since the Δ*codY* mutant in this study grew robustly in nutrient-limited LDM (Fig. 2B), we predicted that it would grow similarly to WT in primary macrophages. We also considered that the Δ*codY* mutant would be deficient in cell-to-cell spread given its stationary phase reduction of ai-BCFAs.

We infected primary bone marrow-derived murine macrophages (BMDM) with Lm strains prepared from mid-log phase LDM cultures and measured viable intracellular bacteria at 0, 4 and 8 hours post infection. At 4 and 8 h post-infection, intracellular growth of the Δ*ilvE* mutant was at least 1 log lower than WT (Fig. 4A). However, the Δ*ilvE* strain showed a growth rate increase after 4 h, suggesting that this strain may be able to adapt to the intracellular environment. The Δ*ilvE::ilvE^+^* strain showed an intermediate phenotype, where intracellular growth was less than that of WT but greater than that of the Δ*ilvE* mutant. WT and Δ*codY* strains replicated within primary BMDM equivalently. We conclude that IlvE is required for optimal growth in the nutrient-limited environment of macrophages, while CodY is not essential for adaptation to intracellular growth within this cell type.

**Figure 4.**
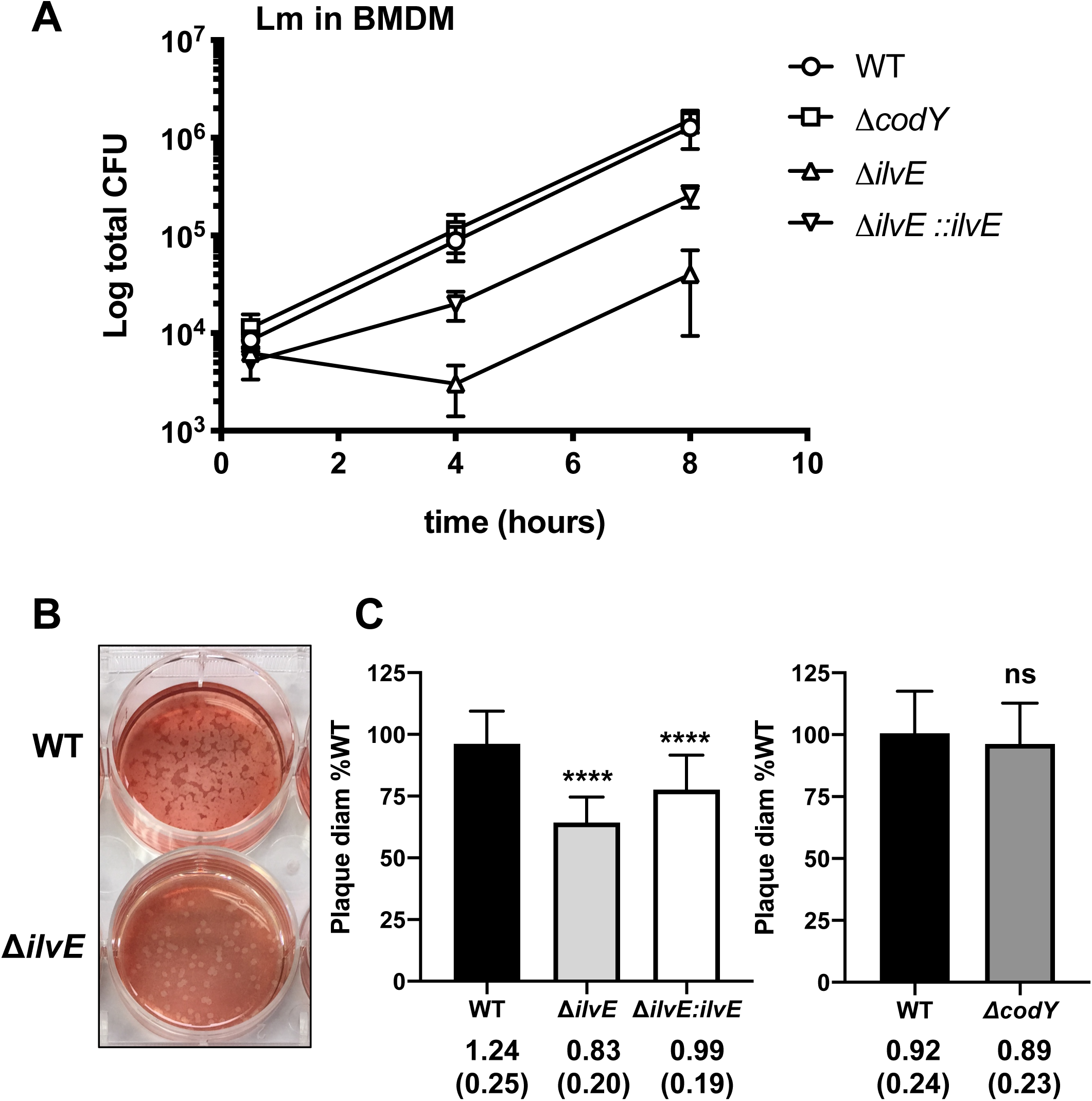
IlvE is required for optimal growth in macrophages and for cell-to-cell spread in cell culture. (A) Total CFU from survival assays of Lm infection of Bone Marrow Derived Macrophages (BMDM) assessed at 0.5, 4 and 8h post-infection. Data are compiled from three independent experiments showing mean and standard deviation. MOI = 1. (B-C) Plaque assay of Lm grown in L9 fibroblasts. (B) Representative image of plaques formed by WT & Δ*ilvE* bacteria after 48h growth. (C) Average plaque diameters from experiments that included WT, Δ*ilvE* and Δ*ilvE::ilvE^+^* (left) or WT and Δ*codY* (right). Numbers below graphs are the mean plaque diameter with standard deviation compiled from three independent experiments. Two-tailed *t-*test comparing mutants to WT, \*\*\*\**P*<0.0001; ns = not significant.

We then infected L929 cells with Lm strains prepared from mid-logarithmic LDM cultures to assess the requirements for IlvE and CodY during multiple stages of intracellular infection as measured by cell-to-cell spread (Fig. 4B-C). Plaques formed from infection with the Δ*ilvE* mutant were approximately 66% the size of WT-infected plaques (Fig. 4C). The complemented Δ*ilvE::ilvE^+^* strain had a partially rescued plaque phenotype. We also observed that plaques formed by the Δ*codY* mutant were not significantly different from those of WT (Fig. 4C). Taken together, these data demonstrate that IlvE is a critical component for Lm intracellular growth and cell-to-cell spread.

### IlvE enhances bacterial survival in response to exogenous membrane stress

Membrane BCFA content underlies Lm resistance to various cell stresses such as pH, small molecules, low temperature, and host-specific antimicrobial mechanisms (8, 10, 12, 13, 16, 17, 35). As a foodborne pathogen, Lm must survive the acidic stomach environment and resist damage from host molecules such as bile. To investigate the role of Ile-dependent BCFA metabolism in protecting Lm membrane integrity, we tested the ability of Δ*ilvE* and Δ*codY* mutants to survive in the presence of membrane disrupting bile salts. We used a bile salt mixture of cholic acid and deoxycholic acid, which are similar to the emulsifying bile acids in the mammalian GI tract. We hypothesized that Lm lacking IlvE would be more susceptible to bile salt stress than WT strains with a full complement of ai-BCFAs. Mid-logarithmic phase bacteria grown in LDM were exposed to 0, 1, 2 and 4 mg/mL bile salts at 37°C for 30 min and measured by counting CFU (Fig. 5A). WT Lm showed decreasing viability with increasing bile salt concentration, with a reduction in viability of almost 2 logs from 0 to 4 mg/mL. The Δ*ilvE* mutant strain showed a consistent 1-log decrease in viability compared to WT at each concentration. The complemented strain Δ*ilvE::ilvE^+^* was slightly less viable at 1 mg/mL, but was similar to WT at 2 and 4 mg/mL. Lastly, the Δ*codY* mutant showed susceptibility to bile salt stress similar to that of WT. We therefore conclude that IlvE promotes resilience against membrane stress, likely through its role in populating the Lm membrane with ai-BCFAs.

**Figure 5.**
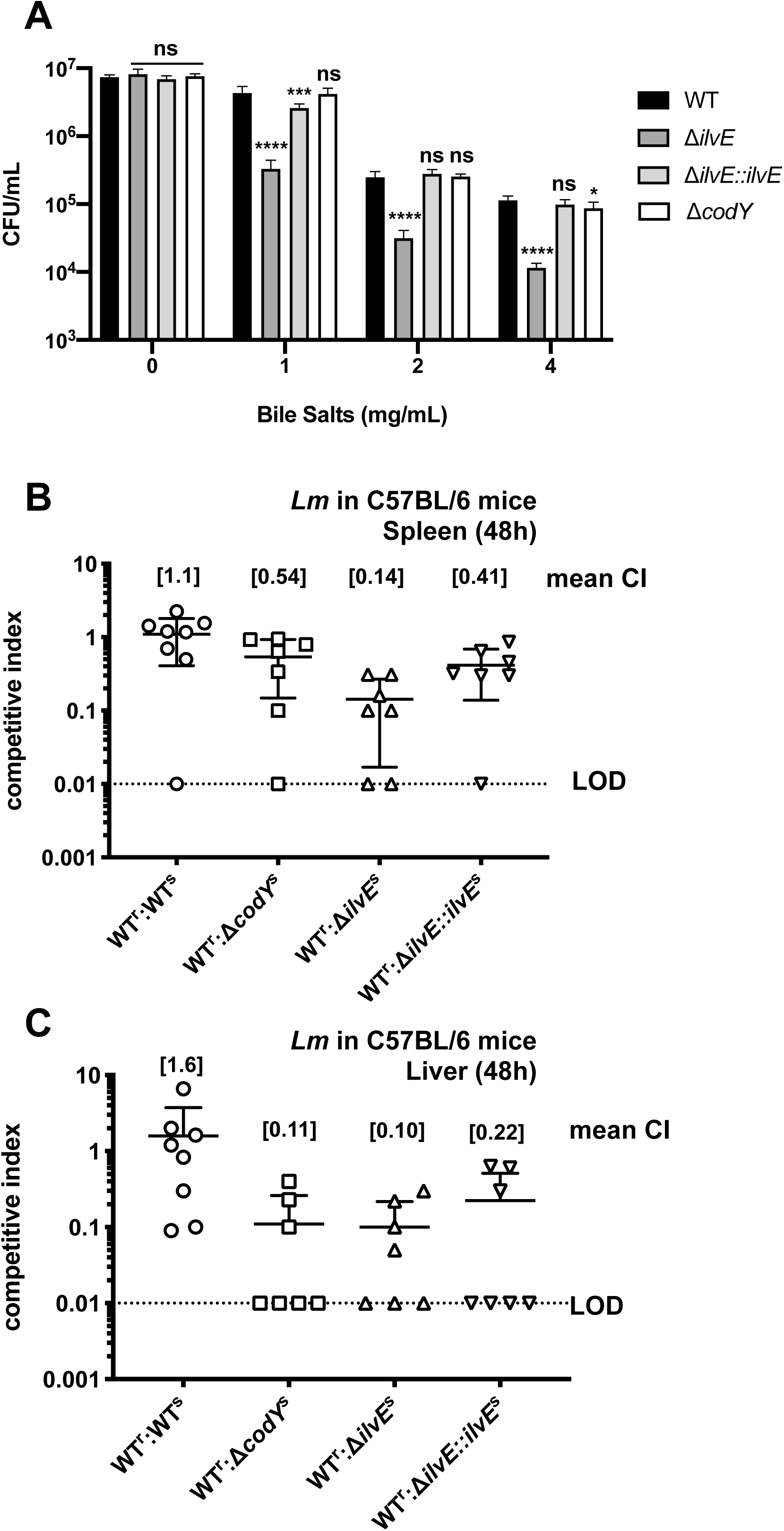
IlvE is required for resistance to membrane stress in response to bile salts and for survival in a mouse model of listeriosis. (A) Log-phase bacteria grown in LDM were added to PBS with 0, 1, 2 and 4 mg/mL Bile Salts (Cholic acid-Deoxycholic acid sodium salt mixture) and incubated at 37°C for 30 minutes. Input for all samples was ∼10^7^ CFU/mL. Data are compiled from three independent experiments. One-way ANOVA (non-parametric) with Dunn’s multiple comparisons post-test comparing mutant strains to WT. ns = not significant; \**P*<0.05; \*\*\**P*<0.001; \*\*\*\**P*<0.0001. (B and C) Female C56BL/6 mice were infected with a 1:1 mixture of erythromycin-sensitive test strains and erythromycin-resistant WT strain via intraperitoneal injection. After 48h infection, (B) spleens and (D) livers were harvested and assessed for viable CFU and competitive index (CI) was calculated as the ratio of Sensitive/Resistant CFU. Data represent two independent experiments with total n=7 mice for all strains except WT, which was n=8. LOD = limit of detection.

### IlvE is required for optimal infection of C57BL/6 mice

While *in vitro* infections can shed light on the intracellular growth capabilities of Lm, they do not illuminate the more complex physiological dynamics of an animal infection. We hypothesized that IlvE and CodY would contribute to pathogenesis in a mouse model of infection, and that the IlvE would have more of an impact due to its constitutive role in membrane fatty acid synthesis. We used a competitive index (CI) assay to measure the fitness of Lm strains in C56BL/6 mice (36). Briefly, we injected mice intraperitoneally with a WT Lm strain that is resistant to erythromycin (WT-erm^r^) combined with a mutant strain (test strain-erm^s^) in a 1:1 mixture (WT-erm^r^ : test strain-erm^s^). After 48 h, spleens and livers were removed and bacteria plated on LB-agar with or without antibiotic to discern resistant (WT^R^) versus sensitive (test strain^S^) bacteria and calculated the CI. The lower the CI, the less “competitive” the test strain was compared to WT during infection.

In both spleen and liver (Fig. 5B-C), substantially fewer Δ*ilvE* bacteria were recovered. The mean CI for the Δ*ilvE* strain in both organs was less than 0.2, indicating severe attenuation in mice. Although the IlvE complemented strain grew better in mice than the deletion mutant, it was recovered at lower levels than WT, suggesting that robust expression of *ilvE* is necessary for optimal survival in a whole animal. Lastly, while bacteria lacking CodY showed a CI of ∼0.5 in mouse spleen, a CI of 0.1 in liver suggests that the liver environment is a more restrictive growth milieu for the Δ*codY* mutant. Overall, these data underline a major role for ai-BCFA metabolism through IlvE for all aspects of Lm growth and virulence, with CodY playing a major role mainly during Ile restriction and severe nutrient restriction.

## DISCUSSION

The plasma membrane of *Listeria monocytogenes* (Lm) is a major structure of the bacterial cell surface and a key interface with host cells (3). Understanding how Lm assembles and remodels membranes to thrive within the host is key to our understanding of this important pathogen. In this study, we explored how two Isoleucine (Ile) responsive proteins, the aminotransferase IlvE and the regulator CodY, contribute to growth, plasma membrane composition, and virulence of Lm. Our findings reveal a crucial role for IlvE in generation of membrane ai-BCFAs, robust growth during nutrient limitation, protection from membrane stress, and virulence in cell culture and in mice. Additionally, our work shows that CodY is involved in modulating membrane ai-BCFA content during stationary phase, and that exogenous Ile is required for bacterial growth when CodY is lacking. However, we observed that CodY is relevant in the nutrient environment of the liver, but contributes less to bacterial fitness in the spleen where Lm primarily replicates in macrophages. Collectively, our findings point to a complex role for Ile usage through IlvE in promoting ai-BCFA membrane composition and also highlight an important relationship between BCAA and ai-BCFA metabolic pathways for Lm pathogenesis.

Anteiso-BCFAs are the major component of the Lm plasma membrane, and the aminotransferase IlvE incorporates Ile into ai-BCFA biosynthesis (Fig. 1) (15, 18). Our main finding that IlvE is a crucial element of Lm biology and virulence is supported first by the observation that bacteria lacking this enzyme (Δ*ilvE*) are severely restricted for growth under multiple conditions of nutrient limitation. Since Lm is an intracellular pathogen, and the intracellular environment is a nutrient-restricted medium (23), Lm must have strategies for acquiring or synthesizing critical metabolites, such as BCAAs, during infection (2, 23). Notably, IlvE was not required for optimal growth in rich-undefined medium, showing that Ile incorporation into ai-BCFAs is not necessary when exogenous nutrients are in great abundance. Rather, IlvE was critically needed for axenic growth during BCAA limitation, in particular when only exogenous Valine or Leucine was unavailable, underscoring the central importance of Ile for membrane metabolism.

Our results also highlight the complex nature of BCAA metabolism in Lm, which is somewhat curious, since these bacteria are able to synthesize BCAA endogenously but still require exogenous BCAAs for optimal growth (22, 26). Lm expresses BCAA biosynthetic genes during infection (26), indicating BCAA limitation within cells. Recent investigation into this phenomenon has revealed that while the Ile-binding regulatory protein CodY inhibits BCAA synthesis when Ile is abundant, the bacteria also limit BCAA synthesis through Rli60 even when CodY inhibition is relieved during Ile restriction, as within the host (21). These opposing processes allow the bacteria to fine-tune Ile levels in order to satisfy BCAA requirements for growth while also allowing virulence gene expression (21). We showed that bacteria lacking CodY or IlvE were severely attenuated for growth when Ile was not available in the medium, highlighting a central role for exogenous Ile during growth. But bacteria lacking CodY harbored more cell-associated BCAAs during growth in nutrient-limited LDM, strongly suggesting a constitutive increase in endogenous BCAA synthesis when CodY inhibition is completely lacking. Thus, the Δ*codY* mutant’s poor growth in the absence of Ile was unexpected, since these bacteria have a greater Ile pool most likely due to *de novo* synthesis. Moreover, the highly robust growth of the Δ*codY* mutant when exogenous Ile and one other BCAA were available underscores a vital role for exogenous Ile in the fine-tuning of BCAA metabolism through CodY, perhaps through involvement in controlling BCAA transport as in *Bacillus subtilis* (27). Collectively, these findings suggest that within the nutrient-restricted intracellular environment, Lm must be able to access sufficient Ile for ai-BCFA synthesis through IlvE activity, but must also sense relative Ile limitation such that CodY metabolic inhibition is relieved to support endogenous BCAA generation for optimal growth.

Another line of evidence pointing to the critical nature of IlvE in Lm biology is its major role in supporting production of resilient membranes during nutrient restriction at biological temperatures (37°C). The importance of ai-BCFA membrane content for resistance to cold has been well established for Lm, and indeed Lm is able to modulate the percentage of ai-BCFAs in response to temperature, salinity, and pH (8, 15–17). However, the Lm membrane is always predominantly made up of Ile-primed odd-numbered ai-BCFAs, emphasizing the central importance of the Ile-to-ai-BCFA biosynthetic pathway for this pathogen. Our demonstration that bacteria lacking IlvE have greatly reduced ai-BCFA content and are sensitive to bile salts directly implicates Ile usage by IlvE as a major player in synthesizing resilient bacterial membranes. Within host cells, Lm is subjected to a variety of membrane-targeting host defenses such as antimicrobial peptides, and ai-BCFAs have been shown to be important for resistance to these mechanisms when the enzyme branched-chain α-keto acid dehydrogenase (BKD), downstream of IlvE, is lacking (13). While those stresses are experienced by Lm inside host cells, Lm is a foodborne pathogen, and so must also survive the low pH of the stomach and the high concentration of bile acids in the small intestine (37, 38). A lifestyle-specific evolution of ai-BCFA metabolism is evident in the Gram-positive dental pathogen *Streptococcus mutans*, which requires IlvE for acid tolerance, such as might be experienced in the oral cavity (39). Thus, the contribution of IlvE for bile salt resistance in Lm reveals that a major need for Ile and ai-BCFAs evolved as a fundamental physiological feature for surviving stress within the diverse environments that this pathogen experiences. Further exploration into the mechanism of bile salt resistance may reveal membrane structural features and bile salt transport mechanisms as playing key roles.

The central importance for IlvE was also revealed by the severe attenuation of the Δ*ilvE* strain in cell culture and in a mouse model of listeriosis. As mentioned previously, the intracellular milieu is nutrient-restricted and a site of antimicrobial assault. Thus, the decrease in intracellular growth of Lm lacking IlvE after four hours of macrophage infection is likely due to enhanced microbial killing, as was seen in the BKD mutant (13). However, it should be noted that the Δ*ilvE* mutant established growth macrophages between 4 and 8 hours, which may indicate a regulatory stress response when Ile incorporation into ai-BCFAs is compromised. This observation, combined with the fact that the Δ*ilvE* mutant still had 20-30% ai-BCFAs during growth in LDM, hints at the presence of another transaminase that can use Ile for ai-BCFA synthesis. Different from what we observed, an Lm mutant lacking *ilvE* in a different parental strain background was almost entirely lacking in ai-BCFAs when grown in rich medium, well under 10% of fatty acid content (14), and this could mean that Lm has several regulatory strategies for membrane homeostasis depending on the nutritional content of the growth medium. However, the amount of ai-BCFAs that we observed in the absence of IlvE was not sufficient for full virulence in a whole animal, highlighting the necessity of IlvE mediated ai-BCFA synthesis for membranes during infection.

Lastly, our results also shed light on the complexity of CodY regulation, which in addition to BCAA metabolism, is also known to be involved in nitrogen and carbon assimilation and regulation of Lm virulence gene expression (21, 25, 27). Previous studies of Δ*codY* mutants in *in vitro* macrophage models have shown different results, where CodY was not required for growth in a transformed macrophage line (25), but was required for optimal growth within primary macrophages (26). In our study, we did not observe a defect in growth within primary macrophages for the Δ*codY* mutant. But note that while the Δ*codY* mutant had an identical fatty acid profile to WT during logarithmic growth, it showed a significant reduction in ai-BCFAs during stationary phase: and for our macrophage experiments, we used Δ*codY* cultures that were prepared at mid-logarithmic phase grown in nutrient-limited medium. This parameter may have poised the bacteria to be more resistant to macrophage killing during the brief, 8-hour duration of the experiment, and this possibility is currently being explored. Regardless, our data are the first to describe a role for CodY in Lm pathogenesis in a whole animal model, where the Δ*codY* mutant was attenuated predominantly in the mouse liver.

In this study, we determined that the branched-chain amino acid transaminase IlvE plays a central role in the membrane dynamics of *L. monocytogenes* and is necessary for robust replication during intracellular infection *in vitro* and *in vivo*. Collectively, our findings highlight an intricate connection between BCAA and BCFA metabolism, and further support a model where Ile is a key metabolite for bacterial growth and virulence, in particular through the activity CodY. Future investigation into how Lm remodels its membrane during interactions with the host will expand our understanding of how pathogens use this defining cellular structure to enhance infection.

## MATERIALS AND METHODS

### Bacteria, cell culture and media

*Listeria monocytogenes* strains used in this study are listed in Supplemental Table S1. Wild Type (WT) *L. monocytogenes* is 10403S and all mutants indicated were created using this parental background. Bacteria were grown in either BHI or LDM (29). Briefly, LDM contains the following final concentrations: 50 mM MOPS/2 mM K_2_HPO_4_ (pH 7.5), 0.02% MgSO_4_*7H_2_O, 0.5 mM Ca(NO_3_)_2_, 0.2% NH_4_Cl, 0.5% Glucose, 0.004% FeCl_3_/Na_3_-Citrate*2H_2_O, 0.5 μg/mL Riboflavin, 1 μg/mL Thiamine-HCl, 0.5μg/mL Biotin, 0.005μg/mL Lipoic Acid, 100 μg/mL of the amino acids Isoleucine, Leucine, Valine, Methionine, Arginine, Histidine-HCl, Cysteine-HCl. Bone marrow derived murine macrophages (BMDM) were isolated from wild type C57BL/6 mice per standard conditions and frozen in liquid nitrogen. The day before *in vitro* infections, cells were thawed, spun by centrifugation, and resuspended in fresh DMEM-10 (Gibco DMEM #11995-065 with 4.5 g/L D-Glucose and 110 mg/L Sodium Pyruvate, 10% Fetal Bovine Serum [HyClone], 1% HEPES [Gibco 1M 15630-080], 1% Non-Essential Amino Acids [Gibco 100X 11140-050] and 1% L-Glutamine [Gibco 200 mM 25030-081]).

### Creation of mutant strains

The markerless, in-frame Δ*ilvE* mutant was constructed using the pKSV7 recombination plasmid (41) per standard conditions such that 1,020 base pairs of the coding sequence were excised. The gene LMRG_02078 sequence in biocyc.org was used for mutant deletion method design. The complemented strain Δ*ilvE::ilvE^+^* was constructed using the Δ*ilvE* parental strain by inserting the coding sequence for LMRG_02078, including 500 base pairs upstream of the start codon, using the shuttle integration vector pPL2 (42) per standard procedures. Note that two independent complemented strains were constructed, one with a FLAG tag inserted at the 5′ end of the gene (Δ*ilvE::ilvE*-FL). Primer sequences are listed in Supplementary Table S2.

The *codY* null mutant was created by insertion-deletion of a *spc* gene originating from the plasmid pJL73 (43). The entire *codY* coding sequence was replaced, in the same orientation, by the spectinomycin resistance cassette using the shuttle vector pMAD (44) per standard procedures. A more detailed description of construction of the *codY* null mutant, including primer sequences, will be included in an upcoming manuscript prepared by T.A. Washington, B. R. Belitsky, and A. L. Sonenshein.

### Growth and survival analysis

Cultures of all strains were grown in liquid LDM or BHI medium to Optical Density 600 nm (OD600) 0.40 – 0.50 and frozen at −80°C in 1 mL aliquots. Frozen stocks were titered for viable bacteria, and on the day of experiments, aliquots were thawed at 37°C for five minutes and shaken at 37°C in fresh medium for 30 minutes. Bacteria were then diluted 1:10 into fresh medium and added to a Bioscreen C honeycomb 100-well plate in a 300 μL volume in triplicate. Plates were incubated at 37°C for 24 hours with constant shaking at medium speed. OD600 readings were taken every 15 minutes on the Bioscreen C instrument. Growth was graphed in Prism. Doubling times were calculated per (40) as follows: n = [log_10_(high OD_600_) – log_10_ (low OD_600_)] / 0.3010 (where OD_600_ values are from exponentially dividing cells). Doubling time = time between OD600 / n.

Survival during exposure to Bile Salts was performed as follows. Strains were thawed from frozen stocks of bacteria grown to mid-log (OD600 ∼0.45) in LDM, added to fresh LDM, and shaken at 37°C for 30 min. Bacteria (∼10^7^ bacteria/mL) were then added to 4 mL of PBS containing Bile Salts (Sigma #48305) at 0, 1, 2 and 4 mg/mL. Tubes were shaken at 37°C for 30 min and then serially diluted with plating on LB-agar plates.

### Fatty Acid Content

Bacteria were grown in LDM to mid-log (OD600 0.4-0.5) and stationary phase (OD600 0.9 – 1.1), spun by centrifugation, washed 1X with PBS, spun again, and frozen at −20°C. Cells were sent on dry ice to Microbial ID for Whole Cell Fatty Acid Analysis. Experiments were performed three times, independently. Results were combined and graphed in Prism 7 or 8 with standard deviation.

### Amino Acid Analysis

Strains grown on BHI agar were used to inoculate fresh liquid LDM and were grown to mid-log (OD600 0.45 – 0.55) or stationary phase (OD600 > 0.8). Cultures (12 or 10 mL) were spun by centrifugation and washed one time with 2 mL 150 mM Ammonium Acetate. Cells were again spun by centrifugation, the supernatant was removed, and cell pellets were snap frozen in a dry ice-ethanol bath. Cells were stored at −80°C until delivery to the Michigan Regional Comprehensive Metabolomics Resource Core (MRC2) at the University of Michigan and analyzed for total amino acid content as follows. Briefly, cells were homogenized in 200 μL of extraction solvent (20% water, 80% 1:1:1 methanol:acetonitrile:acetone) containing 13C or 15N-labeled amino acid internal standards. Samples were incubated at 4°C for 10 min, vortexed, and spun by centrifugation at 4°C for 10 min at 14,000 rpm. Samples were diluted 20-fold and transferred to autosampler vials for mass spectrometric analysis. Chromatographic separation of underivatized amino acids was done using an Intrada Amino Acid column (Imtakt USA). Mobile phases for separation were water:acetonitrile (8:2 v/v) containing 100 mM ammonium formate (solvent A) and acetonitrile with 0.3% formic acid (solvent B). Flow rate was 0.6 ml/min, and sample injection volume was 5 μL. ESI-MS/MS data acquisition was performed in positive ion mode on an Agilent 6410 LC-MS with MRM transitions programmed for both labeled and unlabeled internal standards. A pooled plasma reference sample and “test pooled” sample were included as quality controls. Calibration standards were prepared containing all 20 proteinogenic amino acids at various concentrations and analyzed in replicate along with test samples. LC-MS data were processed using MassHunter Quantitative Analysis software version B.07.00. Amino acids were quantified as pmol/million cells (ascertained by serial dilution and plating) and as pmol/μg total protein using linear calibration curves generated form the standards listed above. All peak areas in samples and calibration standards were first normalized to the peak area of the internal standards.

### In vitro bone marrow derived macrophage infections

Bone marrow derived macrophages (see Bacteria, cell culture and media) were thawed and plated in 24-well tissue culture plates with 2.5 × 10^5^ cells/well and allowed to recover overnight (∼18 hours) at 37°C/5% CO_2_. Following recovery, medium was removed and replaced with 500 μL of DMEM (no antibiotics) containing bacteria (prepped as in Growth Curve analysis) at Multiplicity of Infection (MOI) of one. BMDM with bacteria were incubated for 30 min at 37°C/5% CO_2_ and then washed three times with warm DPBS++ (+Calcium and +Magnesium Chloride – Gibco 14040). One mL fresh DMEM-10 with Gentamicin (50 μg/mL) was added to cells to kill extracellular bacteria. Cells were incubated for 0, 4 and 8 hours. At time of harvest, cells were washed one time with DPBS++ and then incubated in 1 mL of 0.1% Triton-X for 5 min. Cells were removed by scraping and pipetting and then transferred to 3.5 mL sterile double distilled water and vortexed for 10s. 500 μL 10X PBS was added to promote bacterial integrity. Samples were either directly plated or serially diluted and then plated on LB-agar plates and incubated overnight at 37°C. Experiments were done with three technical replicates per experiment on three separate days. Data were compiled and graphed in Prism 7 or 8.

### In vivo mouse experiments

Mouse experiments were performed with 6 to 7-week-old female BALB/c mice. Bacteria were grown in BHI to OD600 0.50 and frozen in 1 mL aliquots. On the day of experiments, bacteria were thawed and resuspended in 3 mL of fresh BHI and incubated with shaking for 1.5 hours at 37°C. Bacteria were pelleted by centrifugation, washed one time with sterile PBS, pelleted again, and then resuspended in 1 mL sterile PBS. Bacteria were serially diluted and plated to ascertain original titer. Bacteria were then combined in the following strain combinations in a 1:1 ratio to attain a concentration of 10^5^ CFU of each strain per 100 μL of PBS. WT-erm^r^:WT-erm^s^; WT-erm^r^:Δ*ilvE-* erm^s^; WT-erm^r^:Δ*ilvE::ilvE^+^-* erm^s^; WT-erm^r^:Δ*codY-* erm^s^. Mice were injected peritoneally with 100 μL of bacterial inoculum. Inocula for all strain combinations were serially diluted and plated on LB-agar and LB-agar-erythromycin plates to measure INPUT concentrations. Mice were then housed for 48 hours in biocontainment rooms before sacrifice and harvest of spleens and livers. Spleens were homogenized in 1 mL sterile PBS with 1.0 mm Zirconia/Silica beads (BioSpec 11079110z), and livers were homogenized in 5 mL sterile PBS with a handheld tissue homogenizer. Samples were serially diluted and plated in duplicate on both non-antibiotic containing LB agar plates and LB agar plates containing erythromycin. CFU/mL per gram of tissue were obtained for all samples sets post-harvest (OUTPUT), and the number of antibiotic sensitive and resistant bacteria were obtained by [CFU on LB plates] minus [CFU on erythromycin-containing plates] = sensitive bacteria. Ratios of erm-sensitive to erm-resistant bacteria for both INPUT and OUTPUT were calculated, and the Competitive index was calculated as OUTPUT ratio / INPUT ratio (36). Mouse experiments were performed on two separate days with n = 3 and n = 4 mice per experiment.

### L929 Plaque Assay

L929 cells (mouse fibroblast cells) were grown in DMEM-10 medium (see “cell culture” above) medium and plated in 6-well tissue culture plates at 10^5^ cells/well at 37°C/5% CO_2_ until cells were almost 100% confluent. On the day of experiments, medium was removed and replaced with fresh medium containing Lm at MOI = 30, incubated for 1h at 37°C/5% CO2, and washed three times with DPBS++ (plus ions). A 1:1 agarose (1.4%):2X DMEM overlay was then added to each well. Plates were incubated at 37°C/5% CO_2_ until plaques were visible. Neutral red mixed with PBS was added to the wells for 1 h to allow for visualization of plaques. After plaques were visible, images of each plate were taken (with a ruler included in the picture), and plaque diameter was measured in ImageJ using the ruler in millimeters (mm) as a standard. At least ten plaques were measured in three separate wells for each of three independent experiments performed on different days. Data were compiled, the mean and standard deviation calculated, and the Student’s unpaired, two-tailed *t*-test was used to compare mutant strains to WT. Data are shown as the mean plaque diameter percentage of WT per each experiment.

### Gene expression via RT-qPCR

Bacteria were grown in LDM to mid-log (OD600 0.4-0.5) and stationary phase (OD600 0.9 – 1.1) and then spun by centrifugation. After lysis by bead-beating, total bacterial RNA was isolated using either the “Quick-RNA Fungal/Bacterial Miniprep” (Zymo Research #R2014) or the “FastRNA Blue Kit” (MPBio #116025-050). RNA was extracted per manufacturers’ protocols and treated with DNase. RNA was precipitated using isopropanol and quantitated on a Nanodrop ND-1000 spectrophotometer. cDNA was made using 250 ng RNA with Invitrogen SuperScript II RT per the manufacturer’s protocol. No-RT controls were created for each RNA sample by omitting RT in cDNA prep. To measure relative gene expression, 1 μL of cDNA was used for SYBR green qPCR using Brilliant II SYBR Green QPCR Master Mix with Low ROX (Agilent #600830) in Bio-Rad Hard Shell PCR 96-well plates (Bio-Rad #64201794) with all cDNA preps done in duplicate, including all no-RT controls. Plates were run on a Bio-Rad CFX96 Real-Time System Thermal Cycler with the following protocol: 10 min at 95°C, 40X cycle of [10s 95°C; 45s 53°C; 1 min 72°C], 30s 95°C, 30s 65°C, 30s 95°C. The *L. monocytogenes* genes *ilvE* (*LMRG_02078*) and *gyrA2* were measured, and *gyrA2* was used as a housekeeping gene for data normalization. Primer sequences are listed in Supplemental Table S2. Changes in gene expression were calculated per the 2^ΔΔCt method (45) comparing mutants to WT.

## ACKNOWLEDGMENTS

This work was funded by NIH R01 AI109048. The authors acknowledge Microbial ID for Fatty Acid analysis and the Michigan Regional Comprehensive Metabolomics Core at the University of Michigan for amino acid analysis. Some Illustrations were partially generated in Biorender (www.biorender.com).

## REFERENCES

1. Freitag NE, Port GC, Miner MD. 2009. Listeria monocytogenes - from saprophyte to intracellular pathogen. Nat Rev Microbiol 7:623–628.

2. Radoshevich L, Cossart P. 2018. Listeria monocytogenes: towards a complete picture of its physiology and pathogenesis. Nat Rev Microbiol 16:32–46.

3. King JE, Roberts IS. 2016. Bacterial Surfaces: Front Lines in Host-Pathogen Interaction. Adv Exp Med Biol 915:129–156.

4. Pizarro-Cerda J, Cossart P. 2018. Listeria monocytogenes: cell biology of invasion and intracellular growth. Microbiol Spectr 6.

5. Carvalho F, Sousa S, Cabanes D. 2014. How Listeria monocytogenes organizes its surface for virulence. Front Cell Infect Microbiol 4:48.

6. Raines LJ, Moss CW, Farshtchi D, Pittman B. 1968. Fatty acids of Listeria monocytogenes. J Bacteriol 96:2175–2177.

7. Kaiser JC, Heinrichs DE. 2018. Branching Out: Alterations in Bacterial Physiology and Virulence Due to Branched-Chain Amino Acid Deprivation. MBio 9.

8. Annous BA, Becker LA, Bayles DO, Labeda DP, Wilkinson BJ. 1997. Critical role of anteiso-C15:0 fatty acid in the growth of Listeria monocytogenes at low temperatures. Appl Environ Microbiol 63:3887–3894.

9. Zhu K, Ding X, Julotok M, Wilkinson BJ. 2005. Exogenous isoleucine and fatty acid shortening ensure the high content of anteiso-C15:0 fatty acid required for low-temperature growth of Listeria monocytogenes. Appl Environ Microbiol 71:8002–8007.

10. Giotis ES, McDowell DA, Blair IS, Wilkinson BJ. 2007. Role of branched-chain fatty acids in pH stress tolerance in Listeria monocytogenes. Appl Environ Microbiol 73:997–1001.

11. Keeney K, Colosi L, Weber W, O’Riordan M. 2009. Generation of branched-chain fatty acids through lipoate-dependent metabolism facilitates intracellular growth of Listeria monocytogenes. J Bacteriol 191:2187–2196.

12. Sun Y, O’Riordan MX. 2010. Branched-chain fatty acids promote Listeria monocytogenes intracellular infection and virulence. Infect Immun 78:4667–4673.

13. Sun Y, Wilkinson BJ, Standiford TJ, Akinbi HT, O’Riordan MX. 2012. Fatty acids regulate stress resistance and virulence factor production for Listeria monocytogenes. J Bacteriol 194:5274–5284.

14. Rogiers G, Kebede BT, Van Loey A, Michiels CW. 2017. Membrane fatty acid composition as a determinant of Listeria monocytogenes sensitivity to trans-cinnamaldehyde. Res Microbiol 168:536–546.

15. Chihib NE, Ribeiro da Silva M, Delattre G, Laroche M, Federighi M. 2003. Different cellular fatty acid pattern behaviours of two strains of Listeria monocytogenes Scott A and CNL 895807 under different temperature and salinity conditions. FEMS Microbiol Lett 218:155–160.

16. Zhu K, Bayles DO, Xiong A, Jayaswal RK, Wilkinson BJ. 2005. Precursor and temperature modulation of fatty acid composition and growth of Listeria monocytogenes cold-sensitive mutants with transposon-interrupted branched-chain alpha-keto acid dehydrogenase. Microbiology 151:615–623.

17. Singh AK, Zhang YM, Zhu K, Subramanian C, Li Z, Jayaswal RK, Gatto C, Rock CO, Wilkinson BJ. 2009. FabH selectivity for anteiso branched-chain fatty acid precursors in low-temperature adaptation in Listeria monocytogenes. FEMS Microbiol Lett 301:188–192.

18. Kaneda T. 1991. Iso- and anteiso-fatty acids in bacteria: biosynthesis, function, and taxonomic significance. Microbiol Rev 55:288–302.

19. Zhang YM, Rock CO. 2008. Membrane lipid homeostasis in bacteria. Nat Rev Microbiol 6:222–233.

20. Eisenreich W, Slaghuis J, Laupitz R, Bussemer J, Stritzker J, Schwarz C, Schwarz R, Dandekar T, Goebel W, Bacher A. 2006. 13C isotopologue perturbation studies of Listeria monocytogenes carbon metabolism and its modulation by the virulence regulator PrfA. Proc Natl Acad Sci U S A 103:2040–2045.

21. Brenner M, Lobel L, Borovok I, Sigal N, Herskovits AA. 2018. Controlled branched-chain amino acids auxotrophy in Listeria monocytogenes allows isoleucine to serve as a host signal and virulence effector. PLoS Genet 14:e1007283.

22. Joseph B, Goebel W. 2007. Life of Listeria monocytogenes in the host cells’ cytosol. Microbes Infect 9:1188–1195.

23. Brown SA, Palmer KL, Whiteley M. 2008. Revisiting the host as a growth medium. Nat Rev Microbiol 6:657–666.

24. Feyaerts J, Rogiers G, Corthouts J, Michiels CW. 2015. Thiol-reactive natural antimicrobials and high pressure treatment synergistically enhance bacterial inactivation. Innovative Food Science & Emerging Technologies 27:26–34.

25. Bennett HJ, Pearce DM, Glenn S, Taylor CM, Kuhn M, Sonenshein AL, Andrew PW, Roberts IS. 2007. Characterization of relA and codY mutants of Listeria monocytogenes: identification of the CodY regulon and its role in virulence. Mol Microbiol 63:1453–1467.

26. Lobel L, Sigal N, Borovok I, Ruppin E, Herskovits AA. 2012. Integrative genomic analysis identifies isoleucine and CodY as regulators of Listeria monocytogenes virulence. PLoS Genet 8:e1002887.

27. Lobel L, Sigal N, Borovok I, Belitsky BR, Sonenshein AL, Herskovits AA. 2015. The metabolic regulator CodY links Listeria monocytogenes metabolism to virulence by directly activating the virulence regulatory gene prfA. Mol Microbiol 95:624–644.

28. Lobel L, Herskovits AA. 2016. Systems Level Analyses Reveal Multiple Regulatory Activities of CodY Controlling Metabolism, Motility and Virulence in Listeria monocytogenes. PLoS Genet 12:e1005870.

29. Belitsky BR. 2014. Role of PdxR in the activation of vitamin B6 biosynthesis in Listeria monocytogenes. Mol Microbiol 92:1113–1128.

30. Biswas R, Sonenshein AL, Belitsky BR. 2020. Genome-wide Identification of Listeria monocytogenes CodY-Binding Sites. Mol Microbiol doi:10.1111/mmi.14449.

31. Brinsmade SR. 2017. CodY, a master integrator of metabolism and virulence in Gram-positive bacteria. Curr Genet 63:417–425.

32. Sonenshein AL. 2005. CodY, a global regulator of stationary phase and virulence in Gram-positive bacteria. Curr Opin Microbiol 8:203–207.

33. Levdikov VM, Blagova E, Colledge VL, Lebedev AA, Williamson DC, Sonenshein AL, Wilkinson AJ. 2009. Structural rearrangement accompanying ligand binding in the GAF domain of CodY from Bacillus subtilis. J Mol Biol 390:1007–1018.

34. Levdikov VM, Blagova E, Young VL, Belitsky BR, Lebedev A, Sonenshein AL, Wilkinson AJ. 2017. Structure of the Branched-chain Amino Acid and GTP-sensing Global Regulator, CodY, from Bacillus subtilis. J Biol Chem 292:2714–2728.

35. Sen S, Sirobhushanam S, Hantak MP, Lawrence P, Brenna JT, Gatto C, Wilkinson BJ. 2015. Short branched-chain C6 carboxylic acids result in increased growth, novel ‘unnatural’ fatty acids and increased membrane fluidity in a Listeria monocytogenes branched-chain fatty acid-deficient mutant. Biochim Biophys Acta 1851:1406–1415.

36. Auerbuch V, Lenz LL, Portnoy DA. 2001. Development of a competitive index assay to evaluate the virulence of Listeria monocytogenes actA mutants during primary and secondary infection of mice. Infect Immun 69:5953–5957.

37. White SJ, McClung DM, Wilson JG, Roberts BN, Donaldson JR. 2015. Influence of pH on bile sensitivity amongst various strains of Listeria monocytogenes under aerobic and anaerobic conditions. J Med Microbiol 64:1287–1296.

38. Begley M, Gahan CG, Hill C. 2005. The interaction between bacteria and bile. FEMS Microbiol Rev 29:625–651.

39. Santiago B, MacGilvray M, Faustoferri RC, Quivey RG, Jr. 2012. The branched-chain amino acid aminotransferase encoded by ilvE is involved in acid tolerance in Streptococcus mutans. J Bacteriol 194:2010–2019.

40. Moat AG. 2002. Microbial Physiology, 4th ed. Wiley-Liss.

41. Smith K, Youngman P. 1992. Use of a new integrational vector to investigate compartment-specific expression of the Bacillus subtilis spoIIM gene. Biochimie 74:705–711.

42. Lauer P, Chow MY, Loessner MJ, Portnoy DA, Calendar R. 2002. Construction, characterization, and use of two Listeria monocytogenes site-specific phage integration vectors. J Bacteriol 184:4177–4186.

43. LeDeaux JR, Grossman AD. 1995. Isolation and characterization of kinC, a gene that encodes a sensor kinase homologous to the sporulation sensor kinases KinA and KinB in Bacillus subtilis. J Bacteriol 177:166–175.

44. Arnaud M, Chastanet A, Debarbouille M. 2004. New vector for efficient allelic replacement in naturally nontransformable, low-GC-content, gram-positive bacteria. Appl Environ Microbiol 70:6887–6891.

45. Livak KJ, Schmittgen TD. 2001. Analysis of relative gene expression data using real-time quantitative PCR and the 2(-Delta Delta C(T)) Method. Methods 25:402–408.

